# Single-cell sequencing unravels the cellular diversity that shapes neuro- and gliogenesis in the fast aging killifish (*N. furzeri*) brain

**DOI:** 10.1101/2021.07.04.450918

**Authors:** Rajagopal Ayana, Caroline Zandecki, Jolien Van houcke, Valerie Mariën, Eve Seuntjens, Lutgarde Arckens

**Author notes:** Corresponding authors: Lutgarde Arckens, Eve Seuntjens. Equal contribution (Rajagopal Ayana, Caroline Zandecki).

## Abstract

The African turquoise killifish combines a short lifespan with spontaneous age-dependent loss of neuroregenerative capacity. The stem cell niches driving neuroregeneration and their molecular signatures remain elusive. To investigate this, we performed scRNA-seq of the adult telencephalon, combined with full-length transcriptomics using ISO-seq. Our results unveil about 25 cell types including neurons and progenitors of glial-and non-glial nature. Subclustering of progenitors identifies four radial glia (RG), and two non-glial progenitor (NGP) cell states. Combining the molecular profiles with spatial mapping of the RG clusters, reveals two spatially divergent astroglia, one ependymal, and one neuroepithelial-like subtype. We propose neuroepithelial-like RG and NGPs to be the start and intercessor populations of both neuro- and gliogenic lineages. Neuronal classification reveals distinct subtypes and lineages corresponding to excitatory and inhibitory neurons. This catalogue of telencephalon cell types is an extensive resource to understand the molecular basis of intrinsic plasticity shaping adult neuro- and gliogenesis.

**Highlights:**

- ScRNA-seq and ISO-seq identified a complete cell catalogue of the adult killifish telencephalon
- Progenitor diversity revealed the presence of spatially-defined (astro)glial subtypes
- A neuroepithelial radial glia population marks the start point of neurogenesis, accompanied by proliferative non-glial progenitors
- Immature neurons form transcriptional subgroups that correspond to excitatory and inhibitory mature neuronal cell types
- This cellular atlas is a basis for studying neurogenesis and neuro-regeneration upon injury, disease and aging

## Introduction

Adult neurogenesis is a complex biological process involving the production and integration of new neurons into the existing neural network. It has been established that postnatal neurogenesis in vertebrates is focused to specific niches in the brain, and the extent of the process varies from species to species ^1,2^. In mammals, neurogenesis is largely confined to two neurogenic zones namely, the sub-granular zone (SGZ) of the hippocampal dentate gyrus and the ventricular-subventricular zone (V-SVZ) of the lateral ventricles ^3,4^ albeit non-canonical sites do exist ^5^. In the dentate gyrus, new neurons integrate in a controlled manner and are functionally relevant for memory and cognition. Many studies have implicated dysfunction of adult hippocampal neurogenesis in an increasing number of brain disorders, such as major depression, and neurodegenerative diseases ^6^. Thus, stem cell and neuroregenerative therapy can be utilized to improve the condition of patients ^7^. The regenerative capacity in the adult mammalian brain is however limited. It is therefore extremely important to elucidate which regenerative strategies are used in vertebrate regeneration-competent animals, like urodeles and fish ^8^. *N. furzeri* has recently gained traction in regenerative medicine since they share more than 90% of orthologous genes with zebrafish, and over 70% with vertebrate genomes ^9,10^. Akin to mammals, the teleost telencephalon is divided into a dorsal pallium and a ventral subpallium. The dorso-lateral pallium is thought to be the equivalent of the mammalian SGZ, while the subpallium is homologous to the mammalian SVZ ^11,12^.

Brain topology and cell types are conserved between mammals and teleost fish. However, the telencephalon in teleosts develops by eversion or outward folding so that the ventricle surrounds the brain parenchyma and the proliferative ventricular zones (VZ) line the border of each hemisphere ^13^. In zebrafish, distinct proliferative zones differ in the rate of development of new cells ^14^, and the apical radial glia (RG) form the primary progenitor population ^15^. Particularly, RG whose cell bodies are located close to the ventricle, have long processes that extend towards the pial surface of the telencephalon, where they terminate on the walls of blood vessels or at the pial surface ^16^. RG occasionally divide and are the equivalent of adult neural stem cells (NSCs) in mouse neurogenic zones. However, rodent GFAP^+^ NSCs are mostly quiescent and give rise to a proliferative, active NSC or a transiently amplifying progenitor type by asymmetric division ^17^. Zebrafish neuroepithelial cells, which are neural progenitor cells inherited from embryogenesis, help in the constant supply of new RG in the VZ that later enter into quiescence and serve as neurogenic NSCs ^18^. Interestingly, previous work on the African turquoise killifish (*Nothobranchius furzeri*) discovered that it is not RG but another highly proliferative population termed non-glial progenitors (NGPs) that actively drives the neuro-differentiation process in the dorsal telencephalon (pallium) ^19^. These NGPs were observed in both juvenile and adult killifish at the apical surface, dispersed in between the RG. They are characterized by a more immature morphology and express markers of division, including PCNA, and neural progenitor markers NESTIN, MSI1, while being devoid of pan-astroglia/RG markers like GLUL, FABP7 or GFAP ^19^. Based on their proliferative character, NGPs could be regarded as similar to mammalian intermediate progenitor cells (IPC) which are known for amplifying the stem cell progeny before differentiating into neurons or glia. However, they seem to be full of self-renewal and neurogenic potential, resembling early neuroepithelial progenitors. Additionally, NGPs have an apical domain contacting the ventricle, and express NESTIN, which are characteristics of neuro-epithelial cells rather than IPC ^19^. Their full molecular signature remains unexplored in adult killifish.

Since the turquoise killifish is fast growing, reaching its full body size in about 6 weeks ^20^, it is of utmost importance to discover which progenitor types support such rapid neurogenesis and attain neuroregenerative ability at young adult age ^10^. This kind of CNS regenerative ability typically exhibited by young teleosts is of high clinical potential, and cannot be easily studied using mammalian models ^21^. As recovery from neuronal loss in injury states is extremely restricted in the case of aged killifish telencephalon ^22^, understanding the proliferative capacity of normal telencephalic niches supported by glial and non-glial progenitors is essential. Therefore, investigating the cellular heterogeneity of the adult brain is a crucial first step to assess how different cells would react to neurorepair in a young vs. aged or diseased context. Equally important are the molecular players that promote neuroplasticity so that it can later be adaptable to neuro-therapeutics.

In this study, we utilized single-cell RNA sequencing to reveal the cellular diversity of the adult killifish telencephalon with a special focus on RG diversity. We could establish transcriptomes of 9616 cells which represent known and novel killifish brain cell types including NGPs, distinctive RG subtypes, immature and mature neurons. Iterative sub-clustering followed by lineage inference analyses of progenitor cells (PCs) and neuronal cells (NC-PCs) separately delineated the neuro- and gliogenic trajectories including four RG subtypes, two NGP types and novel cell types marking the transition between PCs and NCs. The spatial setting for the PCs revealed differential cellular organization in the neurogenic niches. This unique dataset of the adult killifish will be useful for further investigation of cell-type driven mechanisms of brain regeneration upon injury ^10^, aging ^23^ and disease.

## Results

### Single-cell sequencing identifies main cell types in the adult killifish telencephalon

To obtain unbiased knowledge about the different cell types of the adult killifish brain, we performed single-cell RNA sequencing using the 10X Chromium method (Figure 1A). We first optimized the single-cell dissociation protocol and obtained high viability cell suspensions (> 97%) of 6-weeks-old killifish telencephalon (2 samples) ^24^. After sequencing, the sequenced reads were mapped to the killifish reference genome (Nfu_20140502). Mapping statistics showed that only 45% of the single-cell reads confidently mapped to the available *N. furzeri* transcriptome (available through reference gene annotations). To improve the read mapping, we additionally performed Single Molecule, Real-Time (SMRT) Sequencing and Iso-Seq analysis that renders full-length cDNA sequences (no assembly required) to characterize full-length transcripts and isoforms across an entire transcriptome of the 6 weeks adult telencephalon. Thus, we attained better genome and transcriptome coverage and increased the sequenced read mapping from 45 % to 69 % (Suppl. Table 1). We recovered a total of 9616 single cells after integrating samples with an average of 25,800 mean reads per cell and 828 genes per cell (Suppl. Table 1). The two samples clustered well with minimal batch effects upon integration (Figure 1B, Suppl. Figure 1A). Cell quality check and clustering analysis were performed using Seurat. After removing cells with minimum and maximum thresholds for nUMI, nGene, ribosomal and mitochondrial RNA genes, we obtained 9616 high-quality cells (Suppl. Figure 1B-E). The number of principal components or dimensions for further analyses was limited to 30 based on the standard deviation between dimensions (Suppl. Figure 1C). We further performed scaling and normalization, and clustered the cells using dimensionality reduction method, t-distributed stochastic neighbor embedding (TSNE). TSNE-based clustering revealed 23 clusters (resolution = 0.6), of which around 60% were of neuronal origin (Figure 1C-E). Marker genes and all discriminating genes per cluster were identified under the criteria [adj. p-value ≤ 0.05, log_2_FC ≥ 1.2, ≤ −1.2] (Suppl. Table 2). We found five neuronal clusters [MAP2^+^, SYPA^+^], four glial clusters [GLUL^+^, CX43^+^ or FABP7A^+^] and one non-glial cluster (NGP) [PCNA^+^, HMGB2A^+^] (Figure 1D,G, Suppl. table 2). Another small cell population expressed glial as well as neuronal markers and could not be annotated (NA) (Figure 1D). We also identified microglia (MG) [APOEB^+^, AIF1/IBA^+^, LCP^+^], oligodendrocyte precursor cell/oligodendrocyte (OPC/OD) [OLIG1/2^+^, CD9B^+^, SOX10^+^] clusters (, Figure 1D,G, Suppl. Table 2), and three types of vasculature-related cells (Vas1-3; Figure 1D,G, Suppl. Table 2). We also evaluated cell cycle dynamics of all cells and calculated scores for G1/G2M/S phase occupancy based on the cell-cycle associated gene sets. The PCs branch delivered high G2M phase scores, that indirectly mark proliferation in these cells (Figure 1F). The NCs branch displayed a balance in the cell cycle phases indicating differentiated cell states (Figure 1F). Since the killifish is a fairly new animal model, we provide marker genes specific to different cell types present in the telencephalon (Figure 1G, Suppl. Table 2). The top 50 differentially expressed genes per cell cluster marked each population distinctly (Figure 1G). These marker genes in NCs were downregulated in PCs and *vice versa* suggesting that TSNE-based cluster separation was successful. This initial clustering revealed several neuronal, glial and vascular cell types, and showed similarities in cellular build-up between killifish and mammals.

**Figure 1.**
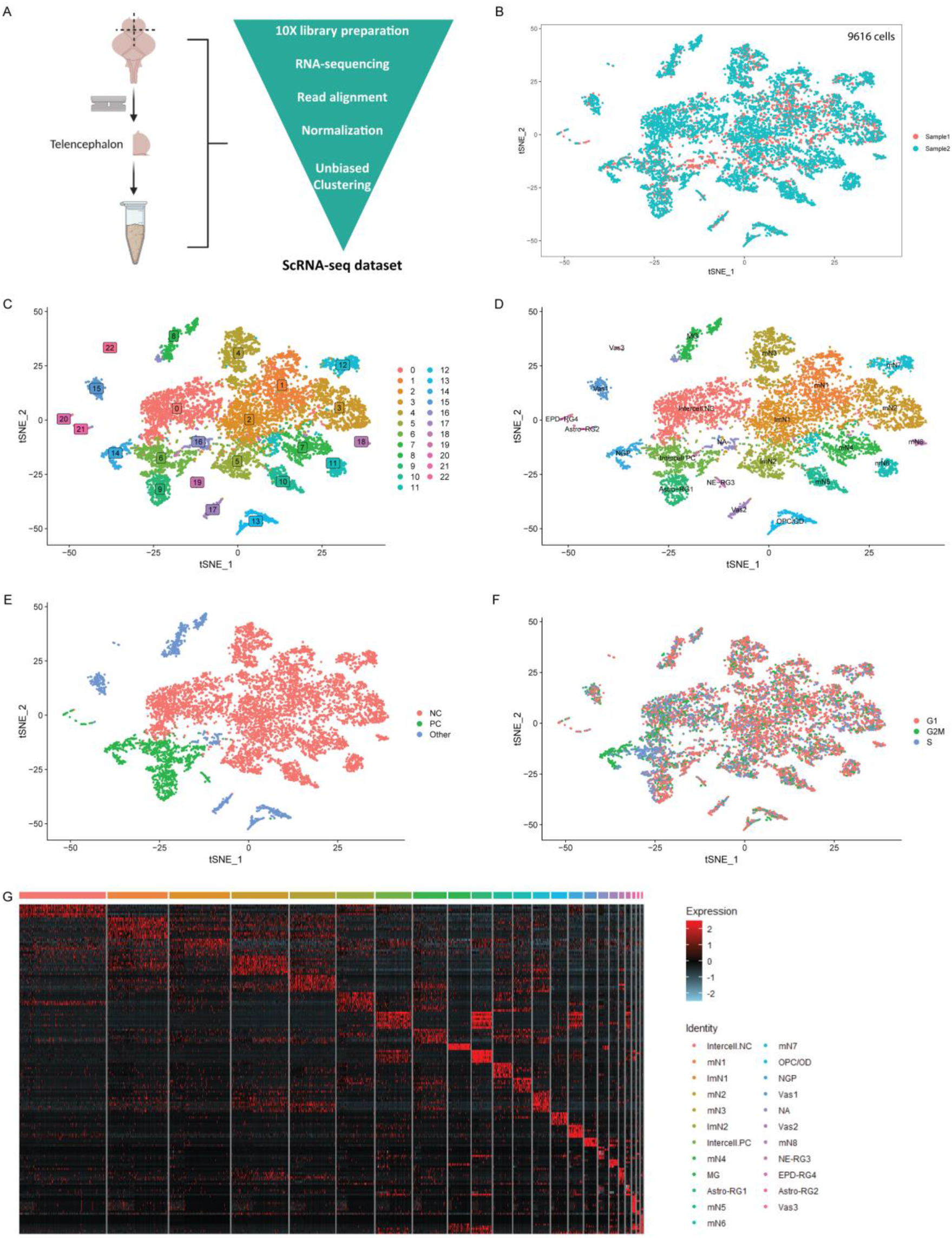
Cell type identification and categorization in the 6-week-old adult killifish telencephalon. **A**) Seurat-based single-cell analysis strategy for 2 samples from the adult whole telencephalon. **B**) Sample-wise overlap in cell numbers revealing minimal batch effects (n = 9616 cells). **C**) TSNE plot at resolution 0.6 revealing 23 cell types comparable to vertebrates. **D**) TSNE Plot with annotation of major cell clusters. **E**) TSNE plot showing the main cell categories, color-coded for NC (Neuronal cells: pink), PC (Progenitor cells: green), Other (All other cells: blue). **F**) TSNE plot showing the Cell cycle phase scoring (G1/G2M/S phases) of individual cells classifying them into proliferative vs. less proliferative cells. **G**) Heatmap of the top 10 markers (ordered by p-value) for the main cell clusters described in D). Note: Relevant gene markers that identify 23 cell types (Suppl. table 2). **Abbreviations**: NGP: non-glial progenitor; RG: Radial Glia; MG: Microglia; mN: Mature neuron; ImN: Immature neuron; Vas: Vasculature; OPC/OD: oligodendrocyte progenitor cell/oligodendrocyte; NA: Not annotated.

### Differential expression of RG subtypes identified in the killifish telencephalon

To obtain a more detailed view on progenitor cell diversity, we sub-clustered all PCs (n = 1303 cells) that expressed markers of division or glial and progenitor markers [PCNA^+^, GLUL(2 OF 2)^+^] (Figure 2A). Sub-clustering unveiled ten cell clusters (resolution 0.8) (Figure 2B), including four distinct RG subtypes and two NGPs. Marker genes and all discriminating genes per cluster were identified under the criteria [adj. p-value ≤ 0.05, log2FC ≥ 1.2, ≤ −1.2] (Figure 2C). The RG subtypes variably expressed FABP7A (BLBP), SEPP1A, SLC1A2B (GLT1), CX43 (GJA1), S100B and GLUL (GS) (Figure 2C-D) which are known teleost RG and mammalian RG or astrocyte marker genes ^25^. Expression of glial marker GLUL was mostly localized to the RG subtypes. Expression of the progenitor marker SOX2 was observed in most PC subtypes (Suppl. Figure 2A, Suppl. table 3) ^26,27^. SLC1A2B and CX43 were used to identify mature astroglia (Astro-RG1, Astro-RG2) in killifish, which have been similarly annotated in mammals, and also juvenile killifish ^19^ (Figure 2C). The density of cells expressing CX43 was similarly high in both astroglial subtypes, whereas CX43 expression was lower or absent in other RG subtypes (Figure 2C-D, Suppl. Figure 2B). Exclusive genes to Astro-RG1 included neurotransmitter transporters SLC1A2B and SLC6A11, metabotropic glutamate receptors GRM3, GRM7 and GABA receptor GABBR1. Astro-RG2 expressed novel genes such as ATP1A2B, ATP1B1, SLC7A3A and astrocyte-enriched markers like SLC3A2, SLC7A5 that are involved in SLC-mediated transmembrane transport ^28,29^ (Suppl. Figure 2B). These characteristics present the first evidence of astroglial diversity in the killifish telencephalon.

**Figure 2.**
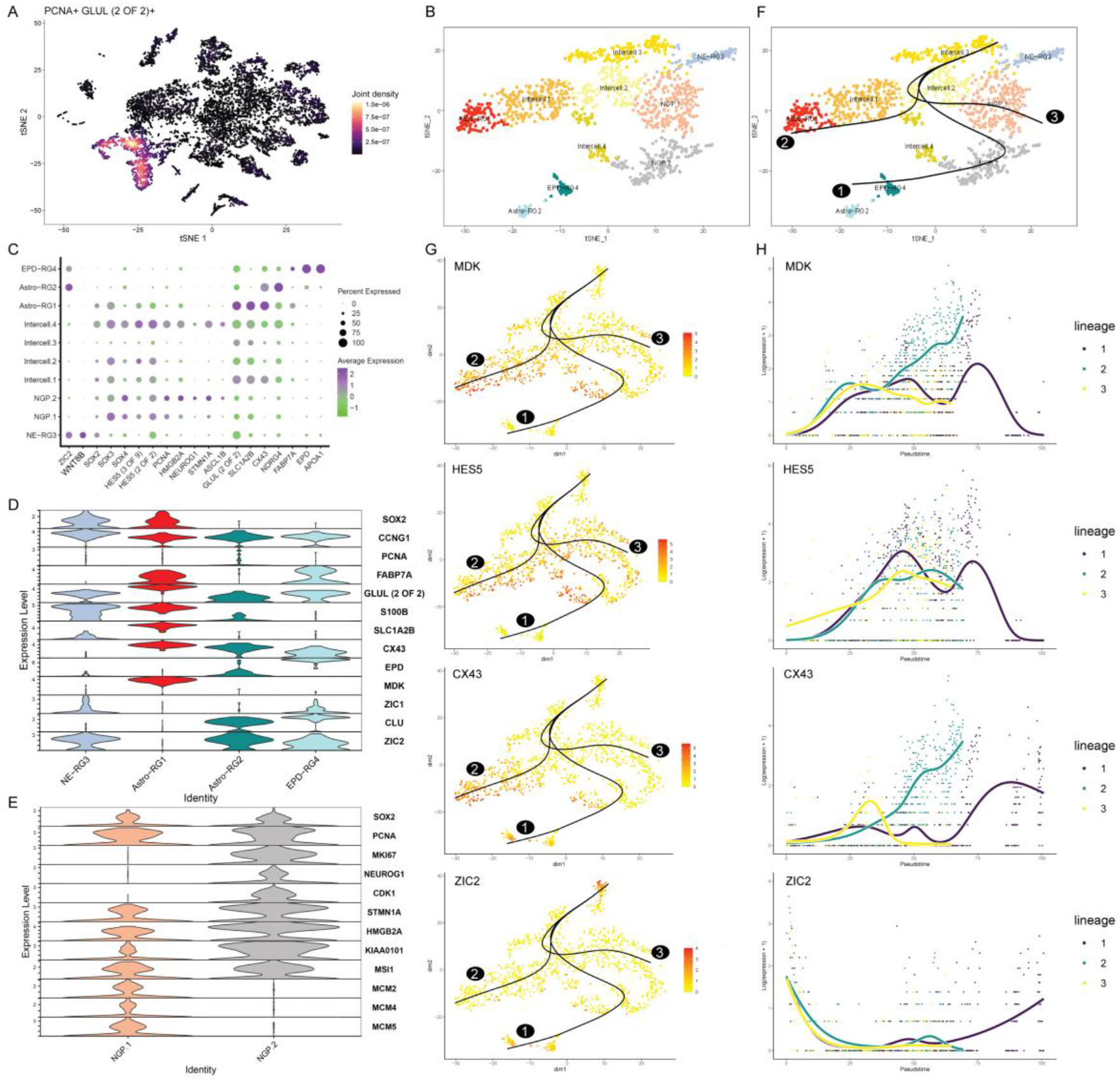
Identification of PCs and relationships. **A)** Strategy for sorting PCs for iterative clustering. **B**) Iterative clustering of PCs shows 10 cell sub-types. **C**) Dot plot shows markers specific to PC sub-types. The size of the dot indicates the percentage of cells expressing the gene (0-100%), the color indicates the expression level. **D**) Discriminating markers of 4 RG sub-types. **E**) Gene expression differences in cell-cycle gene sets between the 2 NGP cell states. **F**) PC-based lineage inference analysis shows 3 divergent trajectories starting with NE-RG3. **G**) Feature plot reveals top differential genes across lineages (scale: yellow to red). **H**) Pseudotime-based plot shows the expression changes across individual lineages in top differential genes. **Abbreviations**: NGP: non-glial progenitor, Inter-cell: Intermediate cells, NE-RG3: Neuroepithelial Radial glia 3, Astro-RG1: Astroglia-like Radial glia 1, Astro-RG2: Astroglia-like Radial glia 2, EPD-RG4: Ependymal Radial glia 4.

Next to Astro-RG1 and Astro-RG2, we identified two other RG types; neuroepithelial RG cells (NE-RG3) and ependymo-glial cells (EPD-RG4). NE-RG3 expressed neuroepithelial cell markers such as ZIC1, ZIC2 and KRT18 (Figure 2D, Suppl. Table 3). Additionally, the expression of RG self-renewal associated factors such as WNT8B, EMX2, SIX3 was also found to be specific to NE-RG3 (Figure 2C, Suppl. Table 3). Due to an everted telencephalon in teleosts, the tela choroidea or ependymal layer surrounds the ventricular surface. In teleosts, the ependyma resembles that of the embryonic mammalian brain with mainly RGs and neuroepithelial cells located at the ventricular surface ^30^ but, in the killifish its structure and function is still a mystery. We found one RG subtype linked to ependyma, EPD-RG4, which expresses FABP7A and ependymin (EPD), and novel markers APOA1, APOA2, APOEB and CLU (APOJ) (Figure 2C-D, Suppl. Table 3). We propose that EPD-RG4 resembles a non-ciliated ependymo-glial cell type as this cluster is devoid of ciliary filament-associated genes ^31^. Thus, these molecular profiles of RGs confirm distinct glial subtypes within the telencephalon and are a first indication of functional and physiological differences.

### Non-glial progenitors account for the bulk of proliferation in the adult telencephalon

An important aspect already explored in killifish larvae recognized RGs as being less proliferative than non-glial cells from 2 days post-hatching (dph) onwards ^19^. In our dataset, sub-clustering of PCs divided the non-glial (NGP) cluster (Figure 1D) into a smaller (NGP.2) and a larger (NGP.1) cluster. These two clusters represent the most proliferative progenitors (PCNA^++^) in the telencephalon (Figure 2C,E). PCNA-associated factor, KIAA0101, HMGB2A and STMN1A were among the marker genes for both NGP.1/2. We analyzed the differences between these 2 populations to ascertain if these are unique cell states or types. While NGP.1 expressed MCM2,4,5 to higher levels, NGP.2 expressed cell cycle genes CDK1,2 and NEUROG1 exclusively (Figure 2E, Suppl. Table 3). It is known that MCM2, 4 and 5 are expressed across all phases of the proliferative cell cycle but are downregulated when cells become quiescent, fully differentiated or go into replicative senescent states ^12^. In NGP.2, pro-neural genes such as ASCL1B, HMGB2B, NEUROG1 ^32,33^, and MARCKS ^34^, were upregulated (Figure 2C,E). HMGB2 is dynamically expressed in proliferating and differentiating NSCs and NPCs ^35,36^. Thus, we hypothesize that the two NGP clusters represent differential cell states with unique molecular programs.

### Four transitional cell states play a central role in gliogenesis

The Intercell.PC population showed a clear skew with a large number of cells scoring high in the S phase (Figure 1F), so we studied this in more depth. In our dataset, sub-clustering of PCs split this Intercell.PC into intermediary sub-clusters [Intercell.1, Intercell.2, Intercell.3, Intercell.4] (Figure 1D, Figure 2C). All Intercell clusters were found to express cellular quiescence-associated markers, HES5 and ID4, which are a part of the Notch and BMP signaling pathways, respectively, and play a vital role in neuro/glial differentiation ^37^, thus supporting a role of these pathways in gliogenesis in the killifish telencephalon (Figure 2C, Suppl. Table 3). These clusters also displayed reduced expression of gene signatures of main PCs (Astro-RG1/2, NE-RG3, EPD-RG4, NGP.1/2), hence suggesting these are cellular states instead of cell types (Figure 2C, Suppl. Table 3). These diverse cell clusters (Figure 2B) that appeared as a single cluster previously (Figure 1D) are clearly separated by the iterative sub-clustering method. The complete list of shared or unique genes between all PC subtypes is available (Suppl. Table 3).

### A dual gliogenic cell trajectory in the telencephalon

Contrary to zebrafish, NGPs and not RG were recently proposed as the neurogenic progenitor type in the juvenile and adult killifish pallium ^19,22,38,39^. Whether the PC subtypes are related to each other and represent different steps in the neurogenic process remains unclear. We performed lineage inference analysis of the PC sub-clusters using Slingshot. The neuro-epithelial cell signature (Figure 2C) and link to adult neurogenesis pose NE-RG3 as the top root cluster candidate (Figure 2F). Analysis using NE-RG3 as the root or start cluster revealed three possible lineages, each of which passed through several Intercell clusters to reach diverse terminal cell states. Using any other cluster as a root yielded physiologically less supported results e.g. terminating in NE-RG3 or NGP.2 (data not shown). The first lineage (1) started with NE-RG3 followed by Intercell.2 to give rise to Astro-RG2 and EPD-RG4 via NGP.2 and Intercell.4, suggesting that EPD-RG4 could be derived from an Astro-RG2 precursor (Figure 2F). This also indicates that at least part of the NGP.2 is gliogenic, for which it requires an additional cell state. The second lineage (2) terminated in the formation of Astro-RG1 via 3 Intercell clusters (Intercell.3, Intercell.2, Intercell.1) (Figure 2F). A third lineage (3) revealed a separate link from NE-RG3 to the NGP.1 cluster via Intercell.3 and Intercell.2. Interestingly, this indicates the presence of independent molecular programs within NGP.1 to possibly become more neurogenic. As the astroglial subtypes (Astro-RG1/2) are formed via different lineages, a definitive functional difference is expected. These lineages (1-2) consistently included Intercell.3/2 as two conserved transition states towards astro-gliogenesis.

We further analyzed genes marking the branches within the lineages via GAM analysis which is used for identifying temporally expressed genes or genes that change with pseudotime. Over the Pseudotime (0 to 100), we assessed the expression of these genes across lineages. The top genes included MDK, HES5, CX43 and ZIC2 (Figure 2G-H). The neural stem cell quiescence marker MDK was lowly expressed initially but increased in lineage 2 rampantly within Intercell.2/3/1 to its peak value in Astro-RG1 (Figure 2G-H). In zebrafish, it has been established that in the telencephalon MDK is mainly restricted to S100B^+^ RG, and in our data, we find correlative expression in S100B^+^ Astro-RG1. HES5, a transcriptional repressor, working downstream of Notch signaling, is known to regulate the timing of neurogenesis and gliogenesis. This gene was expressed heavily in all Intercell clusters. Expression was low in NE-RG3, and increased in lineages 2 and 3 (Figure 2G-H). In case of lineage 1, HES5 expression was more reduced upon reaching maturation in Astro-RG2, which indicates it operates at the time of fate decision, in the Intercell and NGP clusters, in line with its role. As expected, CX43 was found to be lowly expressed in NE-RG3 and NGPs and peaks in terminal clusters Astro-RG1/2, and their possible precursor clusters Intercell.1 and Intercell.4 in lineage 2 and 1, respectively (Figure 2G). We found ZIC2 as a gene upregulated early in pseudotime scale, indicative of an early progenitor marker. ZIC2 was highly expressed in NE-RG3 followed by downregulation across lineages up until re-appearing in Astro-RG2 and EPD-RG4 (Figure 2G-H). In conclusion, trajectory analysis identified a possible neurogenic and two distinct gliogenic routes.

### NE-RG3 have features of multipotent neural progenitors

Our lineage analysis confirmed a root start position for NE-RG3, suggesting they might be the quiescent stem cell population, feeding into the actively proliferating NGP population (PCs lineage 1). Conversely, they could also correspond to a definitive cell cycle exit for terminal glial differentiation as seen in lineage 2 of PCs. To distinguish between these hypotheses, we analyzed the expression of highly conserved molecular players (transcription factors) and pathways of the RG quiescence-to-activation cycle. Transcription factors associated with stemness, such as SOX2, SOX3 and SOX4, were found to be variably expressed in different PCs with relatively high expression of SOX2 in Astro-RG1 and NE-RG3, and SOX3 and SOX4 in NGP.1 (Figure 2C). Cooperation between Notch and BMP pathways is necessary to control stem cell quiescence, and most neural stem cells are known to be quiescent during adulthood ^15,40^. Therefore, we probed for Notch and BMP signaling molecules and downstream effectors such as HES and ID genes within the prominent PC types NE-RG3, NGP.1/2 and Astro-RG1/2 ^41,42^ (Suppl. Figure 2C). In juvenile killifish, it has previously been shown that RGs enter Notch3-dependent quiescence ^19^. In our data, HES1 and HES4 were expressed in NE-RG3, and HES5 in NE-RG3, Astro-RG1 and NGPs. Notch3 expression was specific to Astro-RG1 and NE-RG3; however, higher in Astro-RG1 (Suppl. Figure 2C), thus validating their dormancy. Coolen et al. ^19^ further indicated that NGPs use the pro-neural NEUROG1 in a Notch1-dependent manner and we confirm such similarity to NGP.2, along with expression of Notch1 ligands DLD, DLA (Figure 2C, Suppl. Figure 2C). Within our clusters, ID1 is expressed in most clusters except EPD-RG4, ID2 only in NE-RG3, ID3 in Astro-RG2 and NE-RG3, and ID4 in Astro-RG1 and NE-RG3. What really makes the profile of NE-RG3 reminiscent of quiescent NSCs is the sustained expression of ID1-4, especially ID3 ^37^ (Suppl. Figure 2C). These differential expression profiles indicate the developmental disparity between the different RG cell types (Astro-RG1/Astro-RG2/NE-RG3) found in the killifish ^37^.

As a proxy for quiescence versus proliferative states, we also tested the different PC subtypes using the gene sets specific to different cell cycle phases ^43^. A proportion of cells in the G1 phase was found to steadily increase in the order of NGP.2 < Intercell.4 < NGP.1 < Astroglia < NE-RG3 thus, verifying the most proliferative to the most quiescent cell types (Suppl. Figure 2D). The percent of Intercell clusters in the S-phase was more than NE-RG3 and Astro-RG1, indicative of cycling within the transitioning cells (Suppl. Figure 2D). This reveals that NGP.2 is the most proliferative progenitor cell type, NGP.1 and Intercell.4 come in second, and astroglia (Astro-RG1/2) and NE-RG3 are mostly quiescent (Suppl. Figure 2D). Overall, this supports the theory that NGPs with, possibly, the aid of NE-RG3 are responsible for the explosive telencephalic growth and rapid neurogenesis in the adult killifish ^19^.

### Spatial map of Astroglia and ependyma in killifish

To functionally characterize the two astroglia cell types, we examined their spatial organization via hybridization chain reaction (HCR) for CX43 and SLC1A2B markers (Figure 3). Visualization of mRNA on rostral and caudal coronal sections of the telencephalon revealed a spatially distinct location of Astro-RG1 (CX43^+^, SLC1A2B^+^) and Astro-RG2 (CX43^+^, SLC1A2B^-^) (Figure 3C-D). Astro-RG1 is particularly present at the outer border of the everted telencephalon, lining the ventricular surface. More rostrally, an abundance of Astro-RG1 is detected at the lateral, medial and dorsal surface, almost completely surrounding the two telencephalic hemispheres (Figure 3C, E-H). Caudally, Astro-RG1 is also lining the dorsal and lateral surface (Figure 3D), whereas Astro-RG2 is exclusively present at the ventro-caudal conjunction of the two hemispheres (Figure 3D, I-L). The detection of SLC1A2B and CX43 transcripts in glial fibers extending into the parenchyma further confirms the astroglia-like properties ^44^ of these two RG populations (Figure 3C-D). To characterize the ependymal cells in the killifish telencephalon, HCR targeting EPD was performed (Figure 4A), revealing the distinct presence of EPD-RG4 in the tela choroidea, the thin layer surrounding the ventricular space (Figure 4B-E). The glial gene expression profile (Figure 2C), together with the specific location, identifies EPD-RG4 as a post-mitotic ependymo-glial cell type.

**Figure 3:**
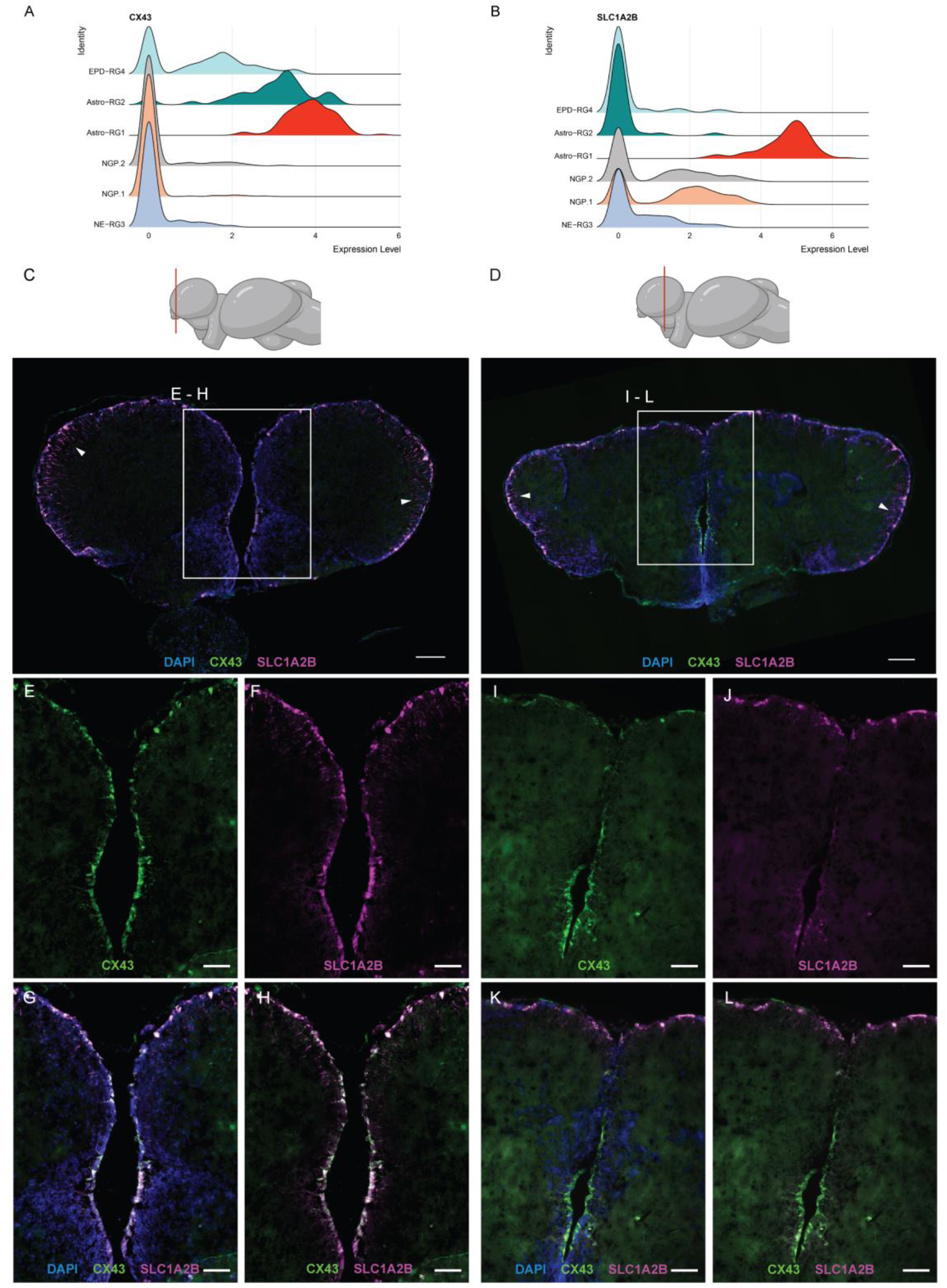
Spatial organization of identified astroglia subtypes Astro-RG1 and Astro-RG2. Ridge plots show the density of cells expressing the astroglial markers **A)** CX43 and **B)** SLC1A2B, in 6 main PC subtypes. The higher the peak, the more cells display a particular expression level in the specific cell type. Multiple peaks indicate differential gene expression within the same cluster. Singular cell peaks at expression 0 are indicative of low/zero expression of the gene in all cells of that cluster. **C**) Rostral and **D**) caudal coronal sections of the telencephalon, the anterior-posterior position of the section is indicated in red on a lateral side view killifish brain illustration. Fluorescent *in situ* labeling of SLC1A2B (magenta) and CX43 (green) mRNA expression in combination with a nuclear stain (DAPI, blue) marks Astro-RG1 and Astro-RG2 respectively. For both sections, CX43 and SLC1A2B expression are present in the cells lining the ventricular surface. Additionally, RG fibers can be identified, extending into the parenchyma, white arrowheads point to zones where these fibers are clearly visible. Scale bar: 100 µm. **E-H)** Magnifications of the zone depicted with a square in the rostral section **C**). Co-labeling of SLC1A2B and CX43 is observed in cells lining the ventricular surface and at the conjunction of the two hemispheres, identifying Astro-RG1 as one of the main cell types at the ventricular surface. Scale bar: 50 µm. **I-L)** Magnifications of the zone depicted with a square in the caudal section **D**). Co-labeling of SLC1A2B and CX43 is observed in cells at the dorsal surface (Astro-RG1). At the conjunction of the two hemispheres, detection of only CX43 mRNA expression is observed, identifying Astro-RG2 as an astroglial-like cell type spatially restricted to the ventro-caudal conjunction of the two hemispheres. Scale bar: 50 µm.

**Figure 4:**
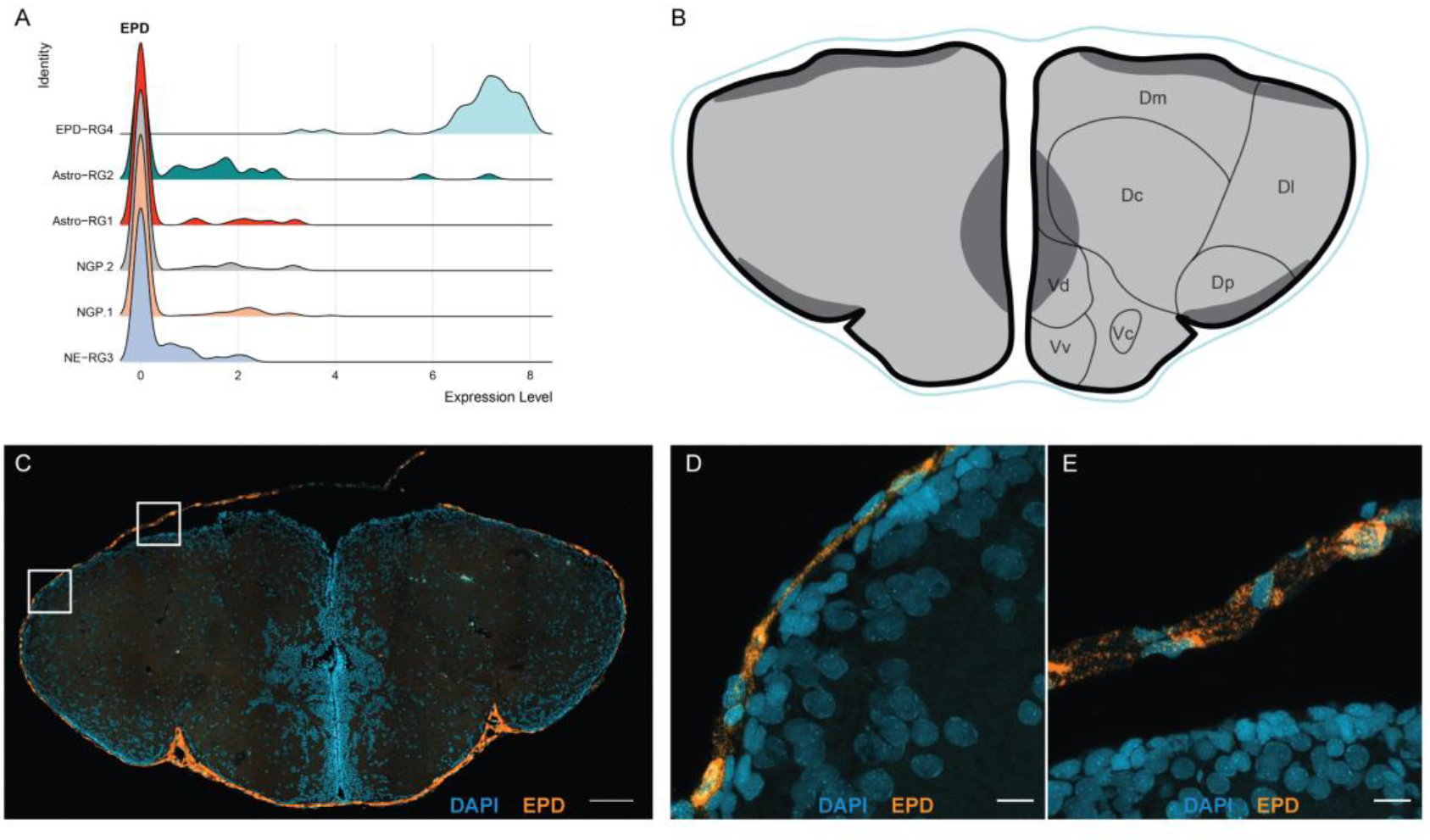
Ependymo-glial cells (EPD-RG4) are found in the tela choroidea. **A**) Ridge plot shows the density of cells expressing the EPD-RG4 marker EPD, in 6 main PC subtypes. **B**) Illustration of the tela choroidea (light blue) surrounding the everted telencephalon as visible on a coronal brain section. The dorsal (D) and ventral (V) subdomains present in the telencephalon are annotated. **C**) Coronal section of the telencephalon showing fluorescent *in situ* labeling of EPD (orange) in combination with a nuclear stain (DAPI, blue). The fluorescent signal can be observed surrounding the ventricular surface. Scale bar: 100 µm. **D-E**) Magnification of zones depicted with squares in **C**). Fluorescent labeling of EPD mRNA expression (orange) is clearly restricted to the tela choroidea. Scale bar: 10 µm. **Abbreviations**: Dc: Central zone of D, Dl: lateral zone of D, Dm: medial zone of D, Dp; posterior zone of D, Vc: central nucleus of V, Vd: dorsal nucleus of V, Vv: ventral nucleus of V.

### Spatial map of starter cell types NE-RG3 and NGPs

High expression of HMGB2A was found exclusively in the NGP.1/2 clusters, making it an excellent marker gene (Figure 5A). STMN1A, known to have an important role in adult neurogenesis, more specifically in the transition from precursor to post-mitotic neurons, was also highly expressed in NGP.1/2 (Figure 5B) ^45^. Visualization of HMGB2A and STMN1A expression together (referred to as “NGPmix”) revealed the presence of proliferating NGPs (PCNA^+^, NGPmix^+^) in all known telencephalic neurogenic niches (Figure 5C, F, G-O) ^46^. A clear difference in the abundance of NGPs could be observed between the neurogenic niches, with region I having a more dense cluster of NGPs at the conjunction of the hemispheres (Fig. 5F-I). On the contrary, at the dorsal and lateral ventricular surface, in region II and III, a more scattered pattern was detected (Figure 5J-R). In these zones, the NGPs were dispersed in between Astro-RG1 (CX43^+^) cells (Figure 5E, J-R). Comparison with gene marker lists for quiescent RGs and proliferative RGs studied in zebrafish ^25^ confirmed that the majority of the RG types in our study were non-dividing [PCNA^--^] (Figure 2C). In contrast, classical glial markers FABP7A, CX43, GLUL and S100B, were enriched in these potentially quiescent or post-mitotic RGs (Figure 2C, Suppl. Table 3). The actively proliferating NGPs thus appeared dispersed with less or non-dividing RG populations in neurogenic regions, as well as grouped into a hotspot of proliferation at ventro-rostral neurogenic region I of the teleost telencephalon ^12,47,48^.

**Figure 5:**
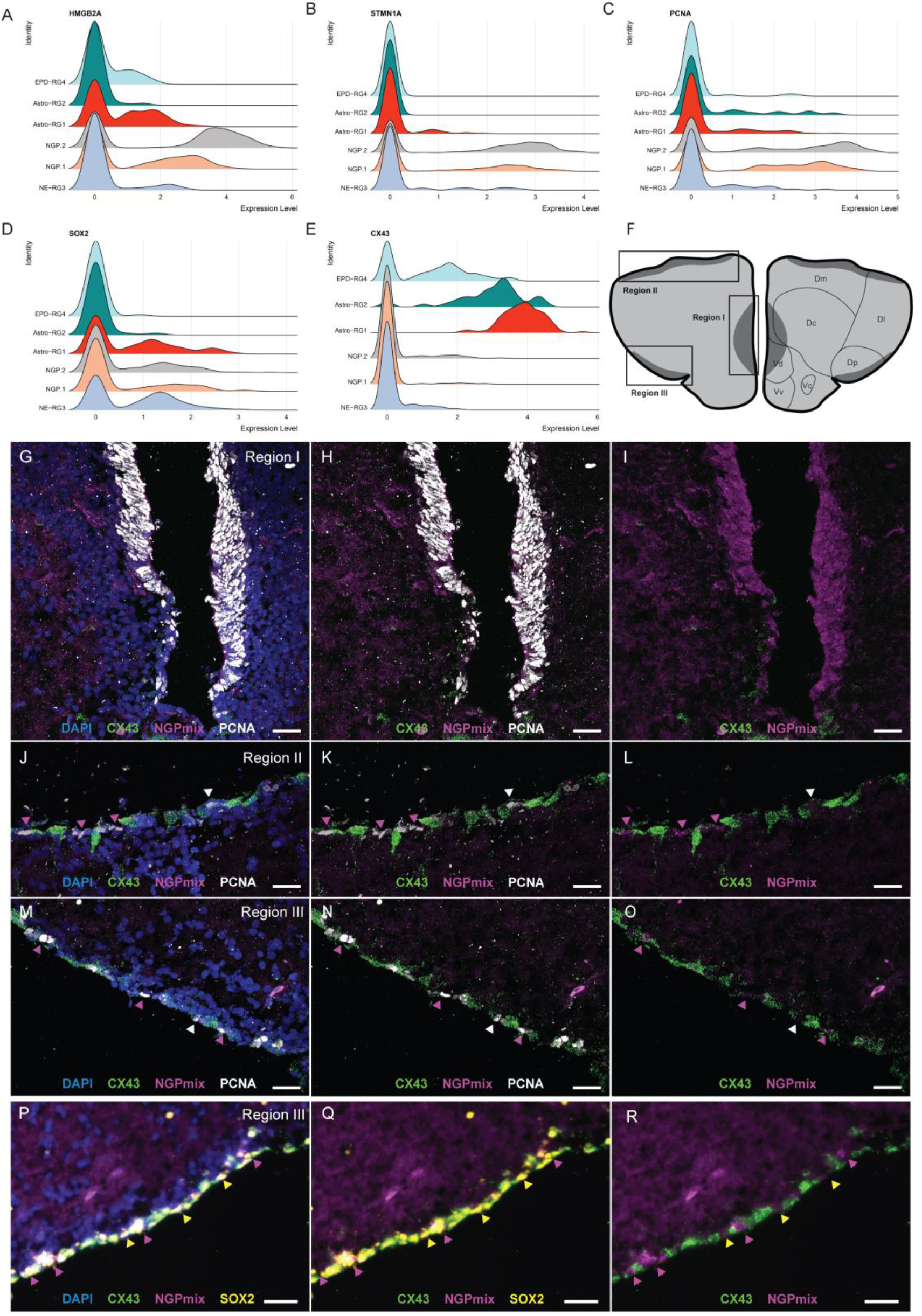
Complementary expression pattern of NGPs and NE-RG3 present in the neurogenic niches. **A-E**) Ridge plots show the density of cells expressing the deterministic markers **A**) HMGB2A **B**) STMN1A, **C**) PCNA **D**) CX43 **E**) SOX2 in main PC subtypes. **F**) Schematic of the 3 neurogenic niches (I-III; dark grey) on a coronal section of the telencephalon. Region I is located at the ventral conjunction of the hemispheres, Region II and III are located at the ventricular surface, at the medial to lateral and posterior zone of the dorsal pallium, respectively. The dorsal (D) and ventral (V) subdomains found in the telencephalon are annotated. **G-O**) Magnification of the 3 neurogenic regions from coronal sections labeled for CX43 (green), HMGB2A and STMN1A (= NGPmix, magenta), in combination with an immune-histochemical staining for PCNA (white) and a nuclear stain (DAPI, blue). In region I (**G-I**), a high abundance of highly proliferative NGPs (NGPmix^+^, PCNA^+^) can be observed, with a limited amount of CX43^+^ RGs at the ventral site of this zone. In region II **(J-L)** and region III **(M-O)**, the NGPs are positioned in between Astro-RG1 (CX43^+^) at the ventricular surface. Magenta arrowheads point to the NGPs (PCNA^+^, NGPmix^+^). White arrowheads point to a limited amount of PCNA^+^, NGPmix^-^, CX43^-^ cells in between the NGPs and Astro-RG1 (green), identified as a small subgroup of dividing RG3 (PCNA^+^ only). Scale bar: 20 µm **(P-R)** Magnifications of neurogenic region III on a coronal section showing fluorescent *in situ* labeling of CX43 (green), HMGB2A and STMN1A (= NGPmix, magenta) in combination with an immuno-histochemical staining for SOX2 (yellow) and a nuclear stain (DAPI, blue): NE-RG3, (SOX2^+^, NGPmix^-^, CX43^-^; yellow arrowheads) can be found in between NGP (magenta arrowheads) and Astro-RG1 (green). Scale bar: 20 µm. **Abbreviations**: Dc: Central zone of D, Dl: lateral zone of D, Dm: medial zone of D, Dp; posterior zone of D, Vc: central nucleus of V, Vd: dorsal nucleus of V, Vv: ventral nucleus of V.

To strengthen the evidence for the presence of NE-RG3 at the ventricular surface, a triple labeling with SOX2 was performed (Figure 5D, P-R). NE-RG3 cells (SOX2^+^, CX43^-^, NGPmix^-^) appear scattered between Astro-RG1/RG2 and the NGPs in the different neurogenic zones. In addition, a few PCNA^+^, CX43^-^, NGPmix^-^ cells were observed at the ventricular surface (Figure 5J-O), indicating a possible minor group of NE-RG3 cells that are dividing in the young adult telencephalon. Since neither Astro-RG2 nor EPD-RG4 expressed SOX2 or other SOX genes (Figure 5C-D), and Astro-RG1, Astro-RG2 and EPD-RG4 (PCNA^-^) can be excluded as the actively proliferating cell types, NGP.1/2 and NE-RG3 are the most plausible neurogenic PC subtypes of the adult killifish pallium. In summary, the spatial distribution together with the molecular profile pinpoints functional differences between NGPs and RG subtypes: two spatially restricted astrocyte-like subtypes, one neuroepithelial and an ependymal type.

### Iterative clustering of NC-PCs revealed intermediate cells and developing neural subtypes

To unravel the ontogeny and relationships between root clusters (NE-RG3 & NGPs) in neurogenesis, we sub-clustered NCs (Figure 1E), Intercell.PC and NGPs (Figure 1D), thus filtering out other glial clusters and non-neuronal cells. Clustering analyses revealed 22 clusters, including prior progenitor types and immature/mature neuron sub-types, with the major contributing populations being a huge diversity of mature neurons (mN, Figure 6A-B). We could annotate nine immature neuronal clusters (ImN [NEUROD2^+^, DCX^+^]), three mature inhibitory neuronal clusters (Ih-mN1/mN2/mN3 [GAD1B^+^ GAD2^+^]) and five mature excitatory neuronal clusters (Ex-mN1/mN2/mN3/mN4/mN5 [SV2A^+^, SLC17A6B^+^]) (Figure 6A-B). The original NGP cluster (Figure 1D) split into two clusters (NGP.1 and NGP.2) and, the Intercell.NC (Figure 1D) split into a larger ImN1 cluster and a smaller Intercell.NC named this way due to its emerging neuronal and reduced progenitor properties [ELAVL3^+^ EOMESA^+^, MAP2^-^].

**Figure 6:**
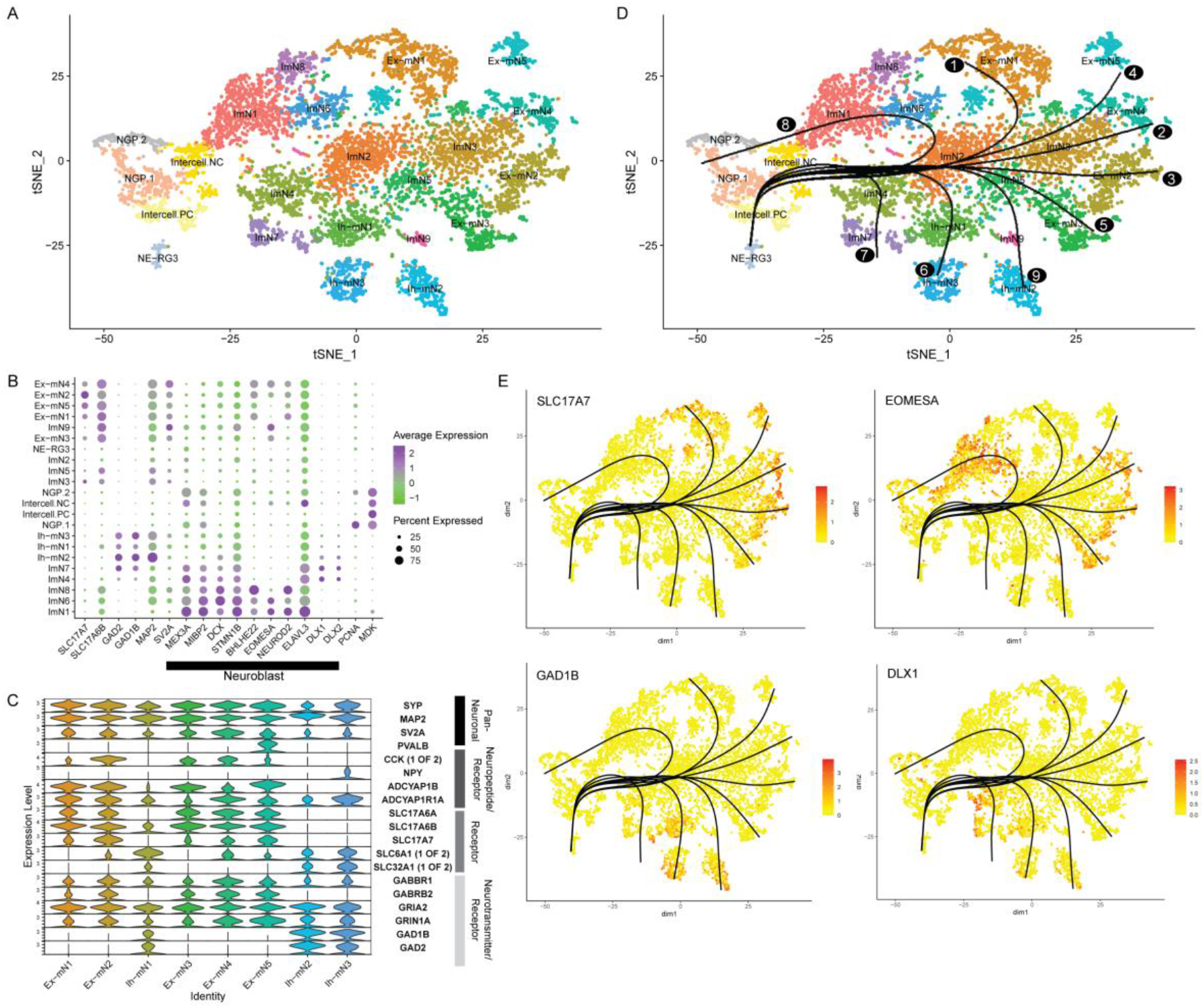
Identification of NC-PCs and relationships. **A**) Sub-clustering of Neuronal clusters (NC-PCs) revealed 22 cell sub-clusters. **B**) Dot plot shows markers specific to all NC-PC subtypes. The size of the dot indicates the percentage of cells expressing the gene (0-100%), color indicates the expression level. Markers of neural progenitor identity and neuronal differentiation have been highlighted in black. **C**) Discriminating markers of 8 mature neuron sub-types and categories. **D**) Pseudotime ordering of cells shows 9 possible lineages (Lineage 1-9) starting from the NE-RG3 cell cluster. **E**) Feature plot reveals top differential genes across lineages. **Abbreviations**: Intercell-NC: Intermediate NC, Ex-mN: Excitatory mature neuron, Ih-mN: Inhibitory mature neuron, ImN: immature neuron, NGP.1/2: non-glial progenitor, Intercell-PC: Intermediate PC, NE-RG3: Neuroepithelial Radial glia 3

MEX3A is required for neuroblast proliferation during killifish adult neurogenesis ^49,50^. In NCs, we found MEX3A expression to increase from NGP.1/2 to Intercell.NC, followed by peak expression in immature neuron subtypes ImN1 and ImN4 indicating the transition from pre-neuroblast to neuroblasts (Figure 6B). MIBP2 is known to be highly expressed in neuroblasts in zebrafish [ELAVL3^+^ EOMESA^+^ MIBP2^+^] ^32^, and we found a similar trend in expression from low in NGP.1/2 to high in ImN1, ImN4, ImN6, ImN7 and ImN8 indicating the formation of fully potentiated neuroblasts (Figure 6B). We also checked for several other immature neuron markers such as ELAVL3, BHLHE22, NEUROD2, DCX, STMN1B (expressed in newborn neurons), and EOMESA (expressed in intermediate progenitors in zebrafish). Interestingly, EOMESA expression was found to be specific to ImN1, 6 and 9 (Figure 6B). Thus, we identified subtypes within immature cell populations that are varying in differentiation stages.

### Divergent expression profiles of the excitatory vs. inhibitory neuronal clusters

In the top five genes expressed by each neuronal cell type, several pan-neuronal markers were found to be shared (Suppl. Figure 3, Suppl. Table 4). Gene commonalities between all mature neuronal clusters (Ex-mN1,mN2,mN3,mN4,mN5, Ih-mN,mN2,mN3), included pan-neuronal markers such as MAP2, SYP and SV2A (Figure 6C). MAP2 is known to label the perikarya, dendrites and axon, while SYP and SV2A label the synapse and presynaptic vesicles. We further aimed to classify the neuron types based on the expression of neuropeptides/receptors, SLC transporters and neurotransmitter/enzyme/receptor genes. Mature neuronal clusters Ex-mN1, Ex-mN2, Ex-mN3, Ex-mN4 and Ex-mN5 hosted an array of features resembling excitatory glutamatergic neurons, such as glutamate transporters SLC17A6A/B (vGLUT2) and SLC17A7 (vGLUT1) (Figure 6C). These clusters also expressed neuropeptide ADCYAP and its respective receptor (Figure 6C). Although we checked the expression of several neuropeptide genes, none of the clusters except Ih-mN3 (NPY^+^) and Ex-mN2 (CCK^++^) showed high expression (Figure 6C). Among all the clusters, PVALB expression was found to be specific to Ex-mN5 which is intriguing as PVALB is often associated with inhibitory GABAergic interneurons in mammals (Figure 6C). We annotated three clusters of mature inhibitory neurons namely Ih-mN1, Ih-mN2, Ih-mN3. All three clusters expressed, next to the glutamate receptor (GRIA1), the GABA synthesizing enzymes GAD1B (GAD65) and GAD2 (GAD67), characterizing them as the primary inhibitory GABAergic neuron populations (Figure 6C). These clusters also shared some specific genes associated with GABAergic transmission like SLC6A1, and the GABA receptor gene GABBR1 but not GABRB2 (Figure 6C, Suppl. Table 4). Thus, the divergent expression profiles of the excitatory vs. inhibitory neuronal clusters reveal a snapshot of the full extent of neuronal diversity in the killifish telencephalon.

### NE-RG3 and NGP contribute to the start of the neuronal lineage

To understand the ontogeny of the neuronal lineage and the relationships between neuronal cell types, we performed a second lineage inference analysis using NCs and PCs (NE-RG3 and NGPs). NC-PC based sub-clustering revealed that NE-RG3 is the root cluster, and both NGP subtypes are linked to neurogenesis (NGP.1/.2) (Figure 2F). Unequivocally, NE-RG3 was marked as the root cluster and this identified nine possible lineages that all passed by NGP.1-NGP.2 clusters, thus confirming their status as the chief proliferative neuronal precursor populations. All the lineages also passed by the novel Intercell.NC cluster which marks the start of the neuronal lineages. We found two major branches in the lineages; the excitatory and inhibitory branch (Figure 6D-E). These are also backed up by specific lineage branch-point markers EOMESA and DLX1/2 for excitatory vs. inhibitory lineages, respectively, which were found to be differential at specific branch points in the lineage-based GAM analysis (Figure 6E).

The first set of five lineages (1-5) passes through the PCs and entered NCs via the Intercell.NC and a subset of immature neuron clusters to reach the excitatory neuronal subtypes (Ex-mN1/mN2/mN3/mN4/mN5) (Figure 6D). The second set of two lineages (6-7) passes via NGPs and Intercell.NC and a different subset of immature neuron clusters to form the inhibitory neuronal subtypes (Ih-mN1/mN2/mN3) (Figure 6D). The final set of two lineages (8-9) indicate the possibility of cell state switching between immature neuron populations, wherein the end-points are ImN1,3,4 (8) or 6-8 (9) but this needs to be further validated (Figure 6D). The dynamic expression of DCX and NEUROD2 among other neurogenic markers within these lineages (7-9) also suggests the presence of reservoir cells that can be used for the prevention of brain aging and disease as proposed in mammals (Figure 6B) ^51^. Thus, lineage inference data confirms NGP.1/.2 to be at the root of neurogenesis.

We analyzed genes marking the branch points within the lineages via GAM analysis and over Pseudotime (0 to 100), we assessed the expression of these temporally expressed genes (Figure 6E). The top genes included DLX1, GAD1, SLC17A7, and EOMESA. In the excitatory neuron lineages (1-4), SLC17A7 expression was high mainly towards the formation of Ex-mNs. Temporal expression of EOMESA was evident with peaks in various immature neuron subtypes that further form the Ex-mNs (Figure 6E), thus indicating a diversity within the telencephalic intermediate progenitor-like cells (EOMESA^+^, DCX^+^). Hence, intermediate progenitor-like cells, that are known to replenish neurons in the postnatal brain are identified. In the inhibitory neuron lineages (5-6), the expression of GAD1 rises only upon the formation of the Ih-mN clusters (Figure 6E). These lineages (5-6) are also marked by temporal expression of DLX1 which is a canonical marker for sub-pallial progenitors and promotes interneuron GABA synthesis, synaptogenesis, and dendritogenesis ^52^. Taken together, our data suggest that neurogenesis in the adult killifish (excitatory or inhibitory) occurs via specific PCs that differentiate to form separate sets of immature neurons or reservoir cell states to finally mature into a neuron.

## Discussion

Single-cell sequencing provides an in-depth perspective of cellular heterogeneity in specific tissues, which is imperative to understand the differential activation of distinct molecular pathways in respective cell types. A central tenet of regenerative biology is that biological processes controlling tissue generation during development often control its regeneration as well. Therefore, processes regulating developmental gliogenesis in the CNS are likely to provide critical insights into neurorepair and its influence on homeostasis in the adult CNS. *N. furzeri*, the shortest living lab-bred teleost model harbors constitutive and active neurogenic niches in all brain subdivisions and thus forms an atypical model to approach this question ^53,54^. Here, we succeeded in identifying the different progenitor cell types and revealed their relationships. One of the key findings is the spectacular cellular diversity in this small region of the killifish brain. The adult telencephalon contains at least 25 cell types (including sub-clusters), and we also identified deterministic markers for all PCs that can be a valuable resource to push the significance of the killifish model in addition to established models such as zebrafish and medaka. We found common vertebrate cell types such as neurons, RG/astroglia, microglia, vasculature-associated cells, neuroepithelial cells, ependymo-glial cells along with the specialized NGPs. We gathered evidence that the decision of cells to become glial or neuronal cells is made within these fast-proliferating NGPs as they seem to be the common progenitor to both glial cell types and neurons. The NE-RG3 population seems primarily quiescent, potentially acting as a kind of reservoir to give the brain the possibility to respond swiftly to injury or disease, or to support explosive brain growth in early life ^19^. Since RGs do not seem to support neurogenesis as much as NGPs do, in the killifish telencephalon ^19,22^, they might be contributing mainly to gliogenic functions.

A subset of radial glia in teleosts has been termed astroglia, and form the ancestral homolog of mammalian radial glia and astrocytes combined due to functional diversification ^30^. Lately, also in zebrafish, astroglia have been gaining attention, and only now it has been established that radial glia are not the only proliferative cell type ^55^. Until recently, it was assumed that bonafide astrocytes do not exist in teleosts and the presence of typical protoplasmic astrocytes was solely a mammalian brain characteristic. It has also been theorized that RGs perform the function of astrocytes in teleosts. Chen *et al.* (2020) reversed the theory by imaging astrocyte morphogenesis in the zebrafish spinal cord ^56^. In line with this, our study discovered diversification within glia, a common process in mammalian astroglia which is seen to rampantly alter with age and disease states ^57^. Spatial analysis of the different progenitor cell types gave a clear insight into the organization of the killifish neurogenic niche. EPD-RG4 were confined to the tela choroidea, whilst most of the dividing and non-dividing progenitors were found to follow a salt and pepper pattern within neurogenic niches I, II and II. We found NE-RG3 to be interspersed with NGPs and astroglia in niche I, in line with its role as a versatile quiescent stem cell. Interestingly, Astro-RG1 was confined to the pallial surface while Astro-RG2 was located in the subpallium, indicating a potentially different origin during development. A potential relation between Astro-RG2 and EPD-RG4 because they are part of the same lineage is intriguing, as this is reminiscent of the generation of mammalian ependymal cells from the sub-pallial V-SVZ ^58^. In zebrafish brain, diverse motile ciliated and non-ciliated cell types within ependymal lineages have been discovered. We found that EPD-RG4 do not express typical ciliary component-associated genes, hence further analyses are needed to explore the inherent diversity within the killifish ependyma ^30^.

Previous studies showed that DCX^+^ neuronal pools are located right below the NGPs in the killifish VZ (only 3-4 cell diameters away from the VZ) ^46^. It was shown that these newborn neurons can be formed in the teleost pallium during development and constitutive adult neurogenesis. Our data indicate that these cells could be the different immature neuron subtypes leading to mature neuron subtypes. In zebrafish, newborn neurons are a derivative of RG mother cells via a non-apical (basal) intermediate progenitor, while in killifish, we show that they are derived from NGP.1/2 (present both in the pallium and sub-pallium) and further intermediate progenitor cells (ImNs) ^25^. The expression pattern of EOMESA in neuroblasts and basal intermediate progenitors in zebrafish VZ ^59^ is similar to that in our killifish Intercell-NC and ImNs [DCX^+^/NEUROD2^+^/EOMESA^+^] clusters, respectively. Thus, these NC populations are more reminiscent of the basal intermediate progenitors than the killifish NGPs [PCNA^+^, EOMESA^-^] ^60^.

NC-PC lineage inferences showed that excitatory as well as inhibitory neuronal clusters can derive from a common source following a single trajectory within the PCs, suggesting that the decision that a neuron would be excitatory or inhibitory might not be driven by the spatial distinction of the stem cell pool. In humans, it was recently shown that cortical progenitors can give rise to both excitatory and inhibitory neurons ^61^. In our data, one can appreciate the divergent neuronal lineages evidenced by high expression of branch point determinists such as EOMESA and DLX1 that govern the excitatory vs. inhibitory lineages (Figure 6D-E). This is also in synchrony with the spatial disparity that is discussed in mammals where the embryonic pallium produces glutamatergic excitatory neurons, while the subpallium is the main source of GABAergic inhibitory neurons ^62^. Having a clear view on the cell type diversity, spatial organization and cell type relationships is an important first step in understanding the neurogenic potential of this emerging new animal model. We note that despite the discovery of the major populations of cells in the growing telencephalon, there is almost certainly additional diversity to be explored in future studies. With this study, we set the stage for future work to address how aging, disease or injury influences the neurogenic and gliogenic landscapes.

## Star Methods

### Fish strains

All experiments were performed on adult 6-weeks-old African turquoise killifish (*N. furzeri*), from the inbred strain GRZ-AD. One male was housed with three females in a multilinking Tecniplast ZebTec aquarium system under standardized conditions; temperature 28°C, pH 7, conductivity 600 μs, 12h/12h light/dark cycle, and fed twice a day with Artemia salina (Ocean Nutrition) and mosquito larvae (*Chironomidae*). All experiments were approved by the KU Leuven ethical committee in accordance with the European Communities Council Directive of 20 October 2010 (2010/63/EU).

### Tissue collection and processing

Fish were euthanized in 0.1% buffered tricaine (MS-222, Sigma Aldrich) and perfused with phosphate-buffered saline (PBS) and 4% paraformaldehyde (PFA, 8.18715, Sigma-Aldrich, in PBS) ^22,63^. Brains were carefully dissected and fixed overnight at 4 °C in 4% PFA. Brains were washed three times in PBS and transferred overnight to 30% sucrose in PBS at 4°C. Next, brains were embedded in 30% sucrose, 1.25% agarose in PBS. Using a CM3050s cryostat (Leica), 10 µm-thick coronal sections were cut and collected on SuperFrost Plus Adhesion slides (10149870, Thermo Fisher Scientific). Sections were stored at −20 °C until the start of HCR.

### Hybridization chain reaction

The probe pair pools targeting CX43, SLC1A2, EPD, STMN1A and HMGB2A (Suppl. Table 5-6) were generated and validated as described in detail before ^64,65^. The selected probe pairs were ordered via Integrated DNA Technologies, Inc (IDT). The HCR protocol (HCR v3.0 ^66^) is based on the protocol of Choi *et al.* (2018), adapted for cryosections as described in Van houcke *et al.* ^22^. In case HCR was combined with immunohistochemistry (IHC), the Proteinase K permeabilization step was removed and the rehydration steps were as follows: the sections were washed in PBS-DEPC 3 times and once in TBS (0,3% Triton-X-100 in PBS). Later, they were washed again in PBS, followed by a washing step in 5x SSCT (0.1% Tween-20 in saline-sodium citrate buffer (SSC)). After the hybridization and amplification steps, the slides were washed 3 times in 5x SSCT before proceeding with the IHC protocol.

### Immunostaining after Hybridization Chain Reaction

After HCR, the sections were washed in PBST (0.1% Tween-20 in PBS) before blocking with 20% normal goat serum (S26, Sigma-Aldrich) in Tris-NaCl blocking buffer (TNB) for 2 hours at room temperature (RT). After blocking, the sections were incubated with the primary antibody diluted in TNB or Pierce Immunostain Enhancer (Thermo Fisher Scientific), depending on the antibody (Suppl. Table 7). After an incubation of 24 hours, the sections were washed with PBST and incubated with the secondary antibody in TNB for 2 hours. Finally, the sections were rinsed with PBS followed by nuclear staining with 4’,6-diamidino-2-fenylindool (DAPI, 1:1000 in PBS, Thermo Fisher Scientific) for 30 minutes. Afterward, the sections were mounted with Mowiol (Sigma-Aldrich).

### Imaging

Images were acquired and processed using a Leica DM6 upright microscope and LAS X software (Leica Microsystems). To obtain more detail, confocal microscopy was used (Fluoview FV1000, Olympus and LSM 900 with Airyscan 2, ZEISS). The images were further processed with Fiji and ZEN software (ZEISS) ^67^.

### Single-cell suspension

3 fish were euthanized in 0.1% buffered tricaine (MS-222, Sigma Aldrich). The blood was removed by intra-cardiac perfusion with cold, sterilized PBS ^22,68^. Next, telencephali were extracted and immediately placed into cold DMEM/F12 (Life Tech, Invitrogen). Telencephali were transferred into sterilized, freshly prepared papain solution (250 µL papain (Sigma Aldrich), 100 µL 1% DNase I (DN25, Sigma Aldrich®), 200 µL L-Cysteine (12 mg/ mL, Sigma Aldrich), in 5 mL DMEM/F12 (Life Tech, Invitrogen)) and digested for 10 min at 37°C. Hereafter, telencephali were dissociated by gently triturating with a cut pipet tip and placed back for 10 min at 37 °C. This procedure was repeated until the brain tissue was completely dissociated into single cells. The suspension was then put through a polypropylene strainer (35 µm, Falcon) and 2 mL of ice-cold, sterilized, freshly prepared washing solution (650 µL D-(+)-glucose 45% (G8769, Sigma Aldrich®), 500 µL HEPES (1M, Thermo Fisher), 5 mL FBS (Life Tech, Invitrogen) in 1X DPBS (Life Tech, Invitrogen) was added. Cells were centrifuged for 10 min at 500 g at 4 °C and the supernatant was discarded. Next, debris was removed via the debris removal solution protocol (130-109-398, Miltenyi Biotec) to ensure high viability of the suspension. Finally, cells were resuspended in cold sterilized 0.04% BSA in PBS. Cell viability was measured at the KU Leuven Genomic Core. A viability of 97.4% was retained after the whole procedure. A detailed description of the single-cell suspension protocol is described in Mariën *et al*., 2023 ^24^.

### Telencephalon SMRT-Sequencing

The Iso-Seq method produces full-length transcripts using Single Molecule, Real-Time (SMRT) Sequencing, thus attaining high accuracy with better genome coverage ^69,70^. We pooled 2-3 fish telencephali (6-week-old) and extracted whole RNA using the RNAEasy Micro kit (Qiagen), with blood. Final RNA quality and integrity were assayed using the DNA 12000 kit on Bioanalyzer (Agilent). Further, this was subjected to PACBIO SMRT Sequencing as recommended in Pacbio protocol for Sequel systems. The SMRT sequencing was performed on the PacBio Sequel at the Genomics Core at KU Leuven (Belgium) using the Clontech SMARTer PCR cDNA Synthesis Kit (recommended by Pacbio). Multiple parallel PCR reactions were set after the first strand synthesis employing the PrimeSTAR GXL DNA Polymerase kit. Next, cDNA of a specific length was selected by splitting these PCR reaction products into 2 fractions, purifying one with 1x AMPure PB Beads and the second with 0.4X AMPure PB Beads, and subsequently mixing both purified products in an equimolar fashion. These libraries were constructed using the SMRTbell template prep kit 1.0. They were sequenced employing Sequel Sequencing, and Binding kits 3.0 on a Sequel I platform with SMRTCells V3.0 LR, allowing a movie collection time of 20h. The mRNAs were selected by the presence of poly-A tails, without 5’cap selection, which was taken into account in the subsequent analysis. The reference genome, gene annotation, and GO terms were downloaded from the *N. furzeri* Information Network Genome Browser (NFINgb). We further used the recommended Pacbio ISOSEQ pipeline to analyze the sequencing data with in-house modifications. We further used the GFFCompare tool to create an updated killifish-specific annotation file by merging the *N. furzeri* reference and Iso-Seq derived full length transcriptome from 6-week-old telencephalon and its specific annotations ^71^. This helped us improve our final genomic annotation file and ensured higher mapping accuracy and coverage for performing Seurat-based single-cell sequencing analysis.

### Telencephalon 10X Genomics sequencing

10X Genomics is known to outperform other droplet-based sequencing methods in terms of bead quality and barcode detection efficiency. To prepare the samples for single-cell sequencing using 10X genomics, all cells isolated from telencephali were pooled from 3 killifish. Subsequently, the single-cell suspension was carefully mixed with a reverse transcription mix before loading the cells on the 10X Genomics Chromium system. Library preparation was done using 10X Genomics droplet-based sequencing with the 10X Chromium Single Cell 3’ (v3 Chemistry) reagents kit. The cells were lysed within the droplet and they released polyadenylated RNA bound to the barcoded bead, which was encapsulated with the cell. Following the guidelines of the 10x Genomics user manual, the droplets were directly subjected to reverse transcription, the emulsion was broken, and cDNA was purified using Silane beads. After the amplification of cDNA with 10 cycles, purification and quantification was performed. The 10X Genomics single-cell RNA-sequencing library preparation involving fragmentation, dA-tailing, adapter ligation, and 12-cycle indexing PCR was performed. After quantification, the libraries were sequenced on an Illumina NovaSeq machine using a HighOutput flowcell in paired-end mode (R1: 26 bp; I7: 8 bp; R2: 50 bp), thus generating 100 million fragments.

### Quality check of single-cell sequencing data

Quality testing of the sparse data matrix (containing genes and cell type sequences) yielded 9616 cells. (Suppl. Table 1). Sequencing metrics showed the presence of 25,800 mean reads per cell with an average gene per cell ratio and absolute transcript or unique molecular identifier (UMI) per cell ratio of 828 and 1687, respectively (Suppl. Table 1). Overall, the cells utilized were sequenced at a depth to obtain optimum coverage of the *N. furzeri* genome (87%). The sequenced reads also confidently mapped to 69% of the killifish transcriptome. On average we detected 17,713 unique genes in both samples (9616 cells). Overall, we could capture high-quality reads (∼97.2% valid cell barcodes) and attain average 57.5% sequencing saturation (Supplementary Table 1).

### Single-cell sequencing data analysis

Raw data were processed with the Cell Ranger 3.0 software. First, we built the reference for CellRanger using the ‘mkgtf’ command (default parameters). The killifish genome (GRZ Assembly, 05/2015 ^8^) as well as the latest NCBI gene annotation (NFIN db), in conjugation with the in-house sequenced killifish ISOSEQ annotation files were used to specify the reference in the ‘mkgtf’ command. Specifically, the two gtf files from PACBIO long-reads specific to telencephalon and the NFIN db reference transcripts were merged using the GffCompare tool. This was followed by the ‘count’ command as part of CellRanger, the option of ‘– expect-cells’ was set to 2500 (all other default options). The results included the feature barcode matrices in various formats usable for downstream analysis in R.

### Data Analysis with Seurat

All matrices from CellRanger were read by the Read10X function using Seurat 4.0 package ^72^. Initially, the 2 samples were processed separately before integration. Initial quality check included removal of ribosomal genes (RPS/L genes) and filtration/removal of cells using the following criteria: a) cells having > 5% mitochondrial content, b) cells < 500 and > 2500 unique genes. We further removed genes/features detected in at least 2 cells (min.cells parameter), and filtered the top 2000 highly variable genes for further analysis. The remaining variable cells and genes were used for downstream analysis. Further, we scaled, normalized and checked the variability of the data using the single SCTranform function. Initially, two Seurat SCTransformed objects, adult killifish and zebrafish scSequencing data as a reference (Cosacack et al.) were used to generate a combined object with IntegrateData function for marker gene identification studies. Post optimization, we only used 2 killifish samples as an integrated object (objects) using FindAnchors and IntegrateData functions and further performed PC and NC sub-clustering. We performed unbiased clustering on the top 30 Principal components which has been represented in the form of dimensionality reductive T-distributed Stochastic Neighbor Embedding (TSNE) at optimized perplexity = n/100 ^73^, and Uniform Manifold Approximation and Projection (UMAP) algorithms. PC or NC-based subclustering was performed by sub-setting pertinent clusters from “all cells” object and rerunning the Seurat analysis.

### Cell type Identification

Cell type identification-related plots (Feature plots and dot plots) were generated by Seurat and cell types were determined by the expression of tested marker genes in the laboratory that define specific cell types and known expression in related species (zebrafish) and humans. We also removed a population of cells occurring to contamination ^74^. The marker genes for all resultant cell populations (All cells, PCs, NCs) were calculated by using FindAllMarkers and FindMarkers function with options min.pct = 0.25, thresh.use = 0.25 and test.use = MAST. Using the FindMarkers and FindAllMarkers function, we ascertained the ranking of all differentially expressed marker genes (threshold: PCT ≥ 0.25) per cell cluster, with associated significance value (corrected p-value ≤ 0.05). The same parameters were applied for the sub-clustering analyses.

### Lineage inference analysis with Slingshot

To elucidate the cell lineage on pseudotime, we made a subset of clustered PC cells (n_cells = 1303), and further converted that Seurat object to a Slingshot object via Slingshot ^75^. Primarily, this is done based on the differential gene expression between cell types by constructing a minimum spanning tree (MST) on cells in a reduced-dimensionality space created by independent component analysis (ICA), and orders cells via a PQ tree along the longest path through this tree. The subset of all cells included NGP.1/2, Astro-RG1/2, NE-RG3 and EPD-RG4 in case of PCs. The ‘start.cluster’ was set to ‘NE-RG3’ and multiple lineages in ‘curves’ mode were analyzed. The ‘start.cluster’ was set to ‘NE-RG3’ and multiple lineages in ‘curves’ mode were analyzed. The estimated size factor was set to 0.6 as the data were already normalized by Seurat. The cells were colored according to the colors on the Seurat TSNE plots. We further performed NB-GAM (negative binomial generalized additive model (NB-GAM)) analysis using tradeSeq, an R package that allows analysis of gene expression along trajectories. We used the patternTest function, which implements a statistical method to check whether the smoothed gene expression is equal along pseudotime between two or multiple lineages. We also checked for early drivers of differentiation using the specifying the knots or branch points of interest, and logging in the gene-specific count information using the plotGeneCount and plotSmoothers function. Similarly, Slingshot and GAM analyses was performed for NC-PC sub-clusters (n=8052) with ‘start.cluster’ set to ‘NE-RG3’. In case of NC-PCs, a subset of first object containing all cells included NGP, NE-RG3, Inter-cell.PC and all NCs.

### Data Availability

The 10X sequencing data have been deposited in GEO (http://www.ncbi.nlm.nih.gov/geo/) under BioProject ID PRJNA718608.

## Author Contributions

**R.A.**: Conceptualization, experimental design, bioinformatics analysis, database development, data visualization, writing: draft, review & editing. **C.Z.:** Experimental design, HCR experiments and analysis, data visualization, writing: draft, review & editing. **J.V.H., V.M.**: Single-cell suspension preparation, writing: review & editing. **L.A, E.S.**: Study supervision, conceptualization, experimental design, writing: review & editing. All authors approved submission.

## Acknowledgements

We thank our collaborators of the KU Leuven killifish consortium Prof. Dr. L. Brendonck and Dr. T. Pinceel. The African turquoise killifish eggs used in this study have been named short-lived GRZ-AD Leuven strain now. We thank Rony Van Aerschot, Arnold Van Den Eynde and Simon Buys, and the KU Leuven killifish team for taking care of the fish facility. We also thank Ruth Styfhals, Dr. S. Gilissen and Dr. M. Hennes for timely discussions. We also acknowledge the help of Ine Jacobs in the HCR experiments. We are grateful to the Genomics Core Leuven for both sequencing experiments and for assistance with data analysis.

## Supplementary figures

**Supplementary Figure 1.**
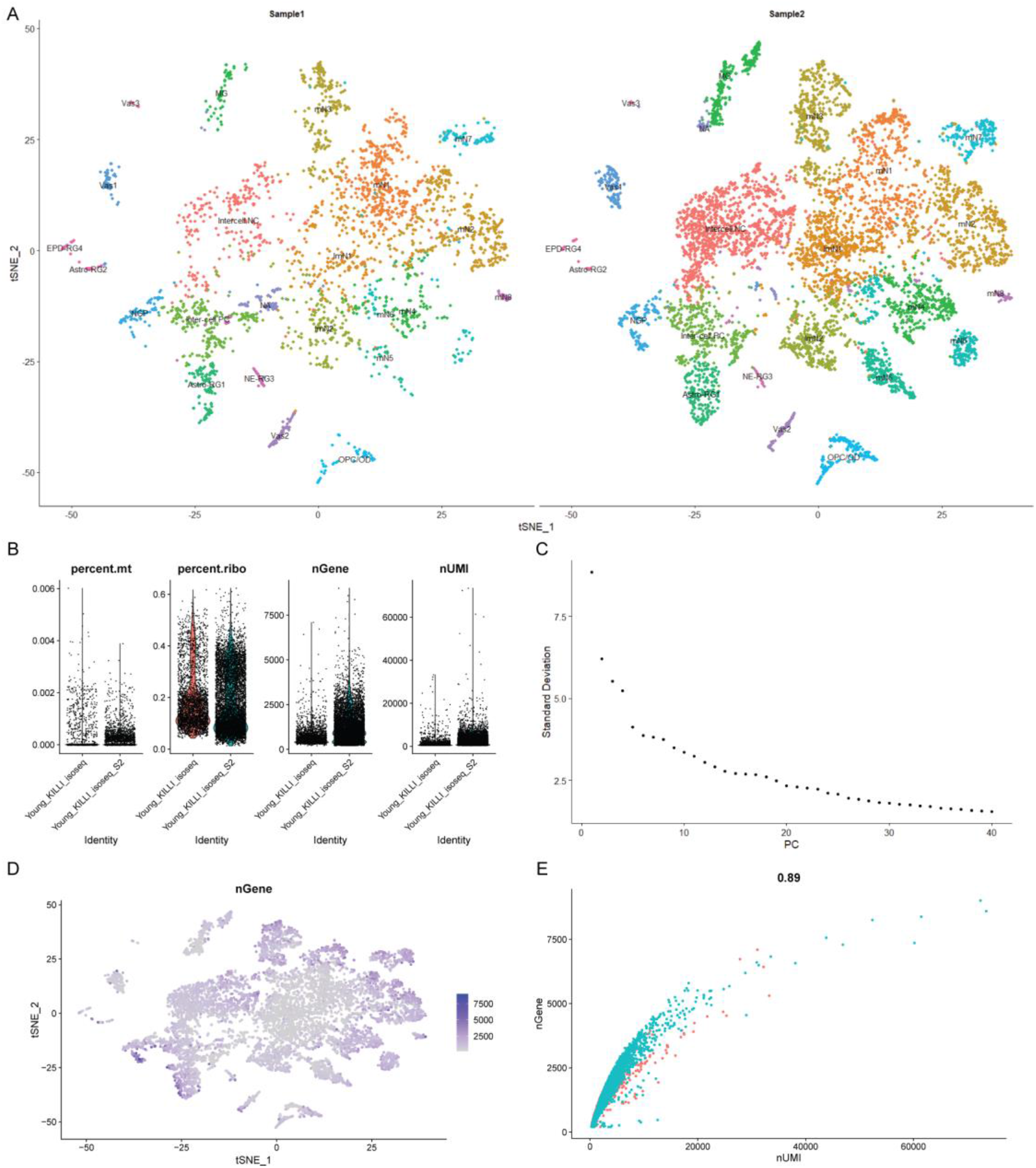
Quality metrics of initial clustering. **A)** Sample-wise overlap of main cell clusters identified in **1D. B**) Violin plots for the number of genes per cell (nGene), number of reads per cell (nUMI), mitochondrial gene percentages (percent.mt) and ribosomal gene percentages (percent.ribo) in nUMI per cell. **C**) Elbow plot shows the most variable principal components among all cells. **D**) Scatter plot showing no. of genes/transcripts detected per cell confirms the high quality of the dataset. **E**) Scatter plot and correlation analysis between nUMI and nGene in all cells.

**Supplementary Figure 2.**
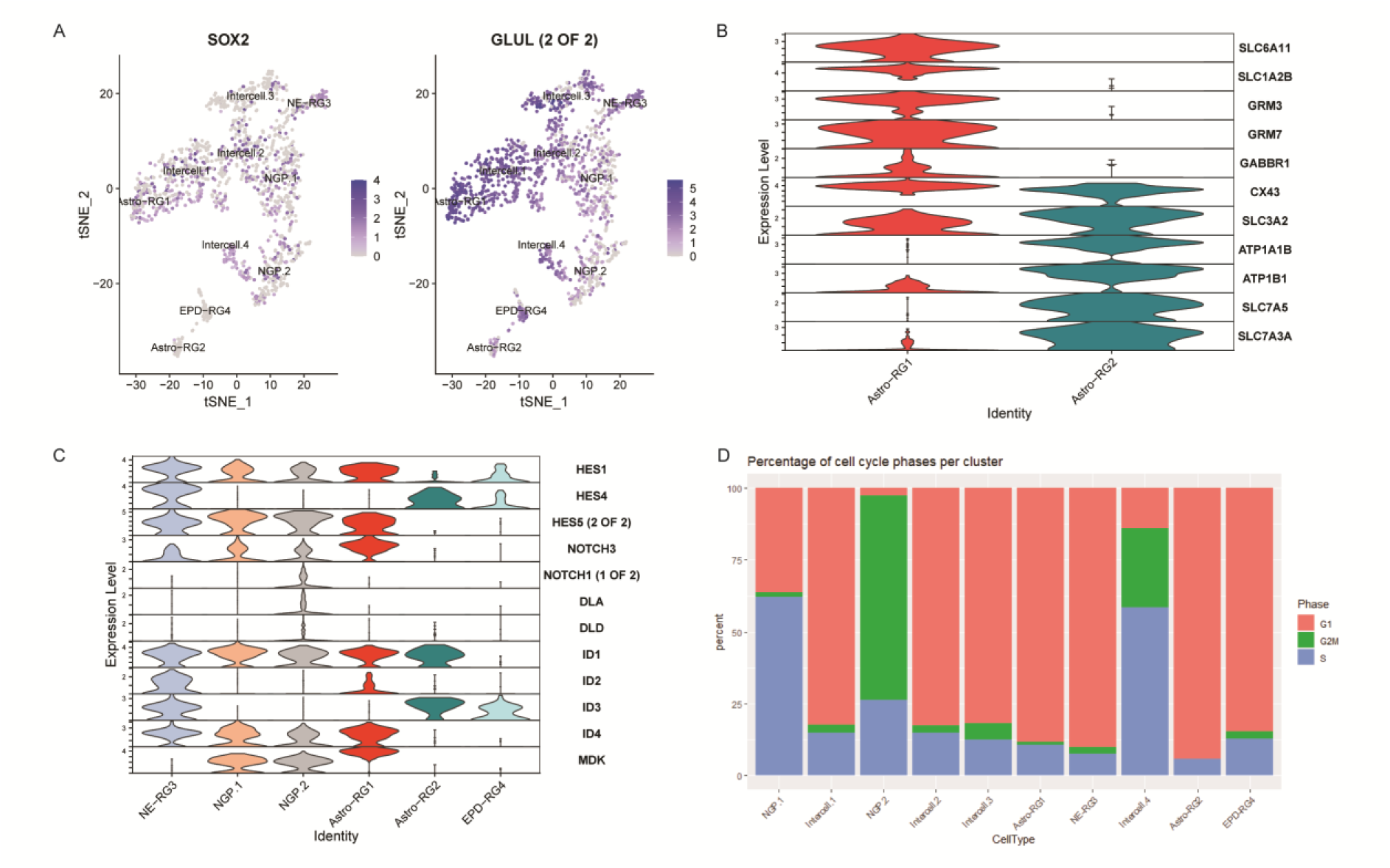
PCs identification and categorization. **A**) Feature scatter plot of all PCs shows the percent of cells expressing SOX2 and GLUL genes. **B**) Discriminating markers of 2 astroglial subtypes. **C**) Violin plot shows the expression of genes associated with HES-ID developmental patterning that govern the cell state gene signatures that promote neurogenesis. Fold changes per gene are mentioned on the y-axis. **D**) Cell cycle phase proportion differences between main PC cell types indicating their level of activity/quiescence: NGPs are actively dividing, whereas NE-RG3 are not.

**Supplementary Figure 3.**
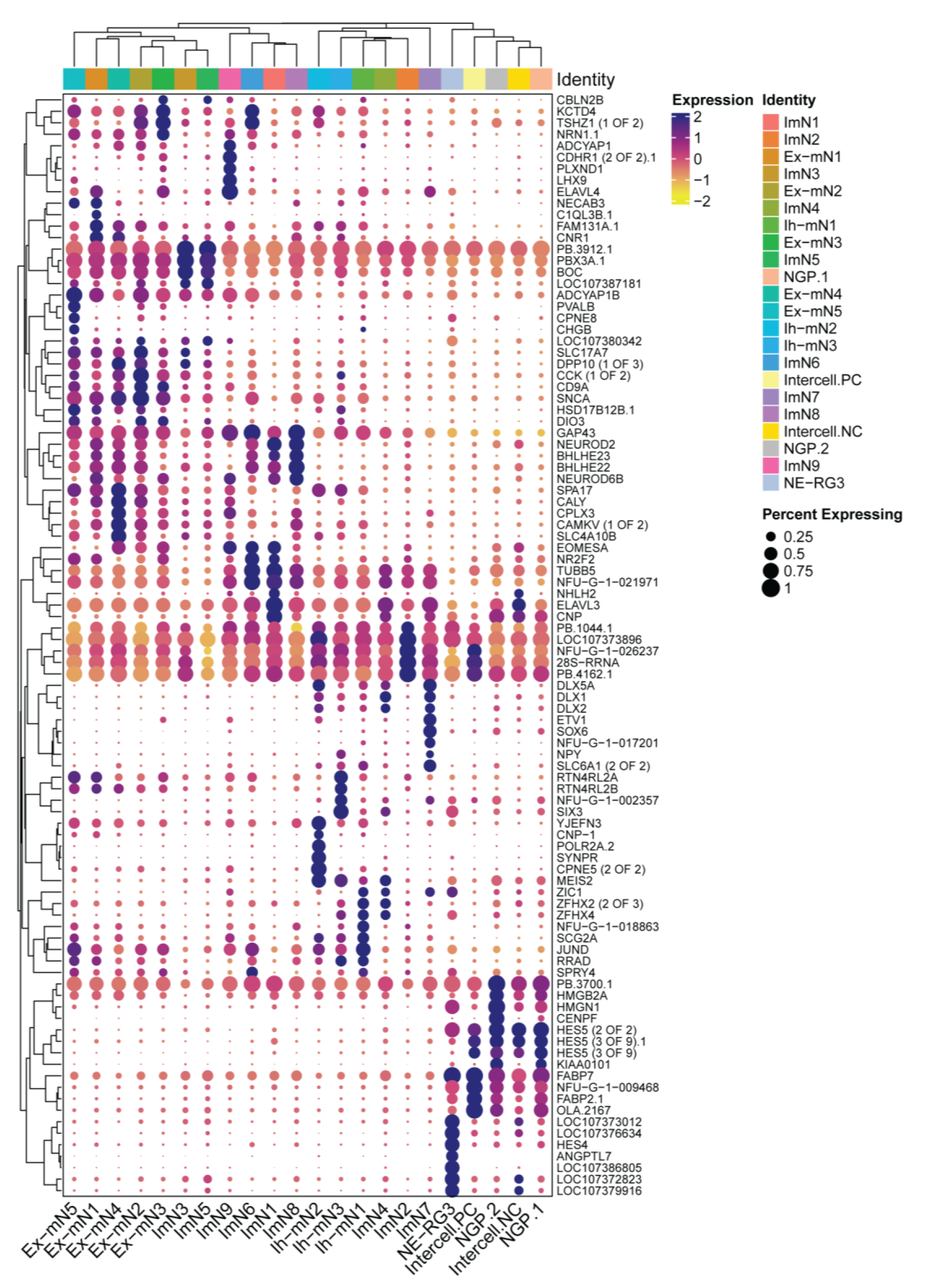
NC-PCs marker details. Top 5 discriminating markers specific to NC and PC sub-clusters.

