## Supplementary table 1-7 for "Single-cell sequencing unravels the cellular diversity that shapes neuro- and gliogenesis in the fast aging killifish (*N. furzeri*) brain": Suppl. Table 6-7.docx

Supplementary Table 6: List of Opools used for hybridisation chain reaction.

| **Opool** | **Target** | **Amplifier** | **No. of probe pairs** |
| --- | --- | --- | --- |
| B3_CX43 | *Nf-cx43* | B3 | 21 |
| B2_CX43 | *Nf-slc1a2* | B2 | 21 |
| B1_SLC1A2 | *Nf-slc1a2* | B1 | 21 |
| B1_STMN1A | *Nf-stmn1a* | B1 | 11 |
| B4_HMGB2A | *Nf-hmgb2a* | B4 | 14 |
| B4_EPD | *Nf-epd* | B4 | 10 |

Supplementary Table 7: List of primary and secondary antibodies used for immunostainings.

| Primary/secondary | Antibody Name | Company | Dilution | Buffer |
| --- | --- | --- | --- | --- |
| Primary | Anti-SOX2, rabbit, SAB2701973 | Sigma-Aldrich | 1:1000 | TNB |
| Primary | PCNA, rabbit, GTX124496 | GeneTex | 1:500 | Pierce Immunostain Enhancer |
| Secondary | Goat anti-Rabbit IgG (H+L) Cross-Adsorbed Secondary Antibody, Alexa Fluor™ 488, A-11008 | Invitrogen | 1:300 | TNB |
